# Pharmacological modulators of epithelial immunity uncovered by synthetic genetic tracing of SARS-CoV-2 infection responses

**DOI:** 10.1101/2022.11.05.515197

**Authors:** Ben Jiang, Matthias Jürgen Schmitt, Ulfert Rand, Carlos Company, Melanie Grossman, Michela Serresi, Luka Cicin-Sain, Gaetano Gargiulo

## Abstract

Epithelial immune responses govern tissue homeostasis and offer drug targets against maladaptation. Here, we report a framework to generate drug discovery-ready reporters of cellular responses to viral infection. We reverse engineered epithelial cell responses to SARS-CoV-2, the viral agent fueling the ongoing COVID-19 pandemic and designed synthetic transcriptional reporters whose molecular logic comprises interferon-α/β/γ-, and NF-κB pathways. Such regulatory potential reflected single-cell data from experimental models to severe COVID-19 patient epithelial cells infected by SARS-CoV-2. SARS-CoV-2, type-I interferons, and RIG-I drive reporter activation. Live-cell-image-based phenotypic drug screens identified JAK inhibitors and DNA damage inducers as antagonistic modulators of epithelial cell response to interferons, RIG-I stimulation, and SARS-CoV-2. Synergistic or antagonistic modulation of the reporter by drugs underscored their similar mechanism of action. Thus, this study describes a tool for dissecting antiviral responses to infection and sterile cues, and a rapid approach to other emerging viruses of public health concern in order to discover rational drug combinations.

## Introduction

Severe acute respiratory syndrome coronavirus 2 (SARS-CoV-2) is the etiological agent of the 2019 coronavirus pandemics (COVID-19), which caused the death or long-term illness of millions of people while simultaneously imposing severe social and economic burden worldwide. COVID-19 is currently managed through massive vaccination, notably with the aid of novel mRNA vaccines ^1^. However, SARS-CoV-2 displayed remarkable evolutionary potential and multiple strains emerged over the last two years, with the Omicron variant currently being the dominant strain globally. Immunity against SARS-CoV-2 strains is limited and continuously threatened by emerging variants ^1^. Pharmacological treatments that reduce the risk of progression in COVID-19 patients are imperfect, and their efficacy is restricted to the early intervention ^2-5^. In particular, the direct intervention through antiviral drugs has been generally underwhelming and the currently approved orally bioavailable prodrug of N4-hydroxycytidine (NHC; Molnupiravir) showed modest efficacy in a placebo-controlled phase II/III trial ^2^, indicating that combination therapies will be required to effectively control disease progression.

One preferential entry site for SARS-CoV-2 is the nasal cavity upon which cells located in the upper respiratory tract, such the nasal epithelial cells and alveolar epithelial type II cells are targeted by the virus. Both share high co-expression of ACE2 and TMPRSS2, the putative main receptor and surface protease respectively, mediating the entry of the several SARS-CoV-2 strains ^6^. SARS-CoV-2 infection leads to transcriptional changes associated with innate immune response in the epithelial cells, including chemokine secretion ^7^, which is believed to trigger a cascade of pro-inflammatory innate immune cell recruitment (macrophages and neutrophils) and disease progression ^7^. Therefore, the first line of defense against SARS-CoV-2 infection is the innate immune system of the upper respiratory tract epithelial cells. One key step of the epithelial cells innate immune response is the production of type I interferons (IFN), which induce an antiviral state in neighboring cells and trigger the activation of the adaptive immune system to protect the host against viral infection. SARS-CoV-2 elicits activation of pattern recognition receptors (PRRs) such as cytoplasmic RIG-I/MDA5 ^8^ as well as TLR2 ^9^ and several other TLRs ^10^. In turn, PRRs activate the nuclear translocation of two families of transcription factors, notably the interferon regulatory factors (IRFs) such as IRF3 and IRF7 and the NF-kB factors p65 and p50, which coordinate the antiviral and acute inflammatory cellular responses. One of the subsequent interconnected events is the transcription of interferon genes, which in turn are secreted and engage the type I interferon receptors (e.g. IFNAR1 and IFNAR2). Interferon activation of the JAK/STAT signaling pathway in both infected and neighbor cells leads to the transcription of interferon-stimulated genes (ISGs) signaling that build up an antiviral state in the cell ^6^. To counteract host cell responses, viruses hijack the activation of the IFN pathway and expression of antiviral genes (aka viral antagonism), limiting the active antiviral response and antiviral signaling ^11^. This leads to the co-existence of viral replication and antiviral response within the infected cell population. Hence, the transcriptional response of epithelial cells is one of the key events that inform on viral infection. Moreover, innate immunity can be activated or amplified by several other pathways, including purely sterile inflammatory environmental triggers such as DNA damage inducers ^12^.

The pathways controlling the transcriptional response of epithelial cells to infection offer an outstanding opportunity for intervention. Whereas drugs with different mechanisms of action targeting the key nodes of the PRRs, NF-kB and JAK/STAT pathways are available and offer the opportunity for creating combination therapies, there is lack in technologies to rationally design combination treatments with synergistic antiviral activity and less prone to resistance. Furthermore, novel drugs and combinations thereof identified through phenotypic drug discovery platforms are potential game changers ^13^, but the phenotypic readouts for SARS-CoV-2 infection and RNA viruses in general are limited. Recently, we developed a method to generate synthetic locus control regions (sLCR) to enable genetic tracing of complex phenotypes, notably transcriptional inflammatory responses ^14,15^. Using an evolved computational approach to design phenotype-specific sLCRs named logical design of synthetic cis-regulatory DNA (LSD; Company C., Schmitt M.J., Dramaretska Y. et al., *manuscript in preparation*), we generate sLCRs designed to intercept the transcriptional response of epithelial cells to SARS-CoV-2 infection using the earliest publicly available signatures in cell models ^16-18^. We characterized one reporter for genetic tracing of cellular response to SARS-CoV-2, which represents cell states including those described in upper airway tracts of severe COVID-19 patients ^7^. This sLCR responds to cellular stimulation by the interferon pathway and of the PRRs. Hence, it is serving as a platform for unbiased drug discovery of synergistic modulators of epithelial immune responses from viral and sterile triggers, and to study their mechanism of action.

## Results

### Designing sLCRs for SARS-CoV-2 productive infection in epithelial cells

To design SARS-CoV-2 reporters that would model infection response in epithelial cells, we first defined the input for LSD, namely SARS-CoV-2 signature and transcription factor genes (**Fig. 1a**). We overlaid bulk and single gene expression data from the earliest available reports in lung epithelial cells over-expressing the ACE2 SARS-CoV-2 entry receptor (A549+ACE2 ^16^) and endogenously expressing ACE2 and the preferential SARS-CoV-2 protease TMPSSR2 (Calu-3 ^16^), as well as from primary gastrointestinal organoids ^17^. In response to SARS-CoV-2 infection, each cell type expressed private and shared genes between 12-60 hours post-infection, and we selected signature and transcription factor genes that were up-regulated in at least two out of five datasets. Pathway enrichment analysis revealed that both of the selected sets are enriched in interferon and acute inflammation response genes (**Fig. 1b-c**). As the selected transcription factors have the highest impact of CREs determination and ranking by LSD, we decided to systematically design SARS-CoV-2 genetic tracing reporters using a combinatorial selection of all transcription factors and signature gene lists. This resulted in 64 potential reporters (**supplementary table S1**)

**Fig. 1.**
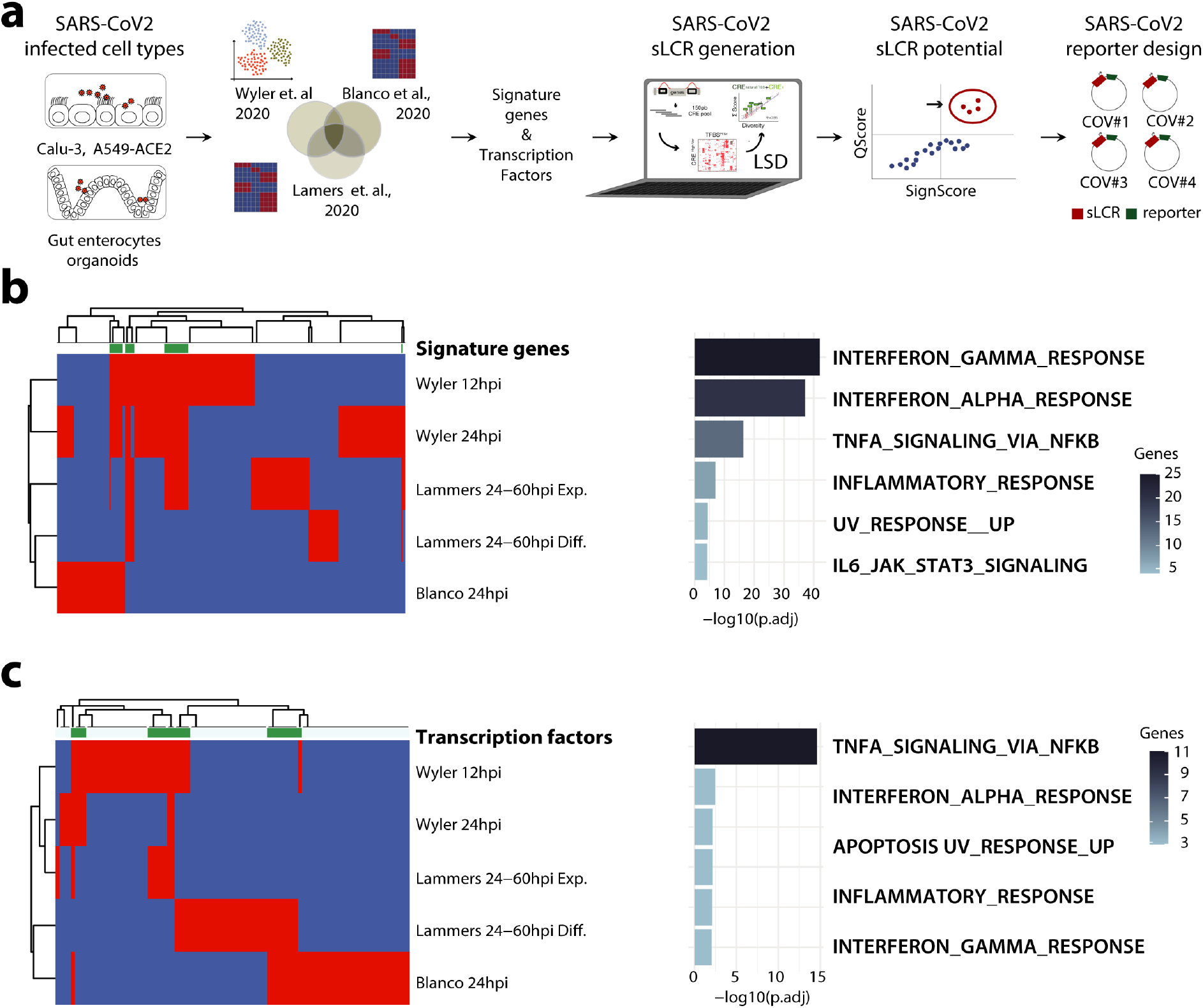
Generation of COVGT sLCRs from defined inputs using the LSD method. Schematic representation of the SARS-CoV-2 sLCR generation. Bulk and single-cell expression profiles from the indicated studies were combined to infer an epithelial response signature to SARS-CoV-2 and define LSD input. **(B-C)** Signature selection to generate the SARS-CoV-2 sLCR. Left, heatmaps of ssGSEA enrichment in the indicated dataset for signature genes **(B)** and transcription factors **(C)**. Green denotes the genes meeting the threshold of activation in >2 different common genes among different datasets. Right, molecular hallmarks enrichment analyses. (Euclidean distance, complete linkage)

To select one of the above-designed reporters that specifically marks productively infected single-cells, we next made use of our recently developed phenotypic ranking approach (**methods**; Company C., Schmitt M.J., Dramaretska Y. et al., *manuscript in preparation*). First, we aimed at specificity towards SARS-2. To this end^19^, we clustered naive, SARS-CoV, and SARS-CoV-2 single-cells from parallel scRNA-seq experiments in one cell type, upon comparative dimension reduction and found two cell clusters that appear specific for the response to viral reads accumulation (C11 and C13), with C11 being specific for SARS-CoV-2 (**Fig. 2a**). Next, we used our TFBS enrichment-based phenotypic potential analysis to rank all our reporters (Company, Schmitt, Dramaretska et al., *manuscript in preparation*) and found that four sLCRs bear a homogeneous scoring higher than other specific or background reporters when both C11 and C13 were assessed (**Fig. S1**). Strikingly, our synthetic reporters COVGT1-to-4 markedly outperformed the phenotypic potential of endogenous interferons used in broadly available reporters (e.g. Addgene #102597, #17596, #30536 and #17598), indicating that synthetic *cis*-regulatory elements have higher on-target potential than endogenous promoters of the human interferon gamma or beta genes, which are expected to respond to SARS-CoV-2 infection (**Fig. S1**).

**Fig. 2.**
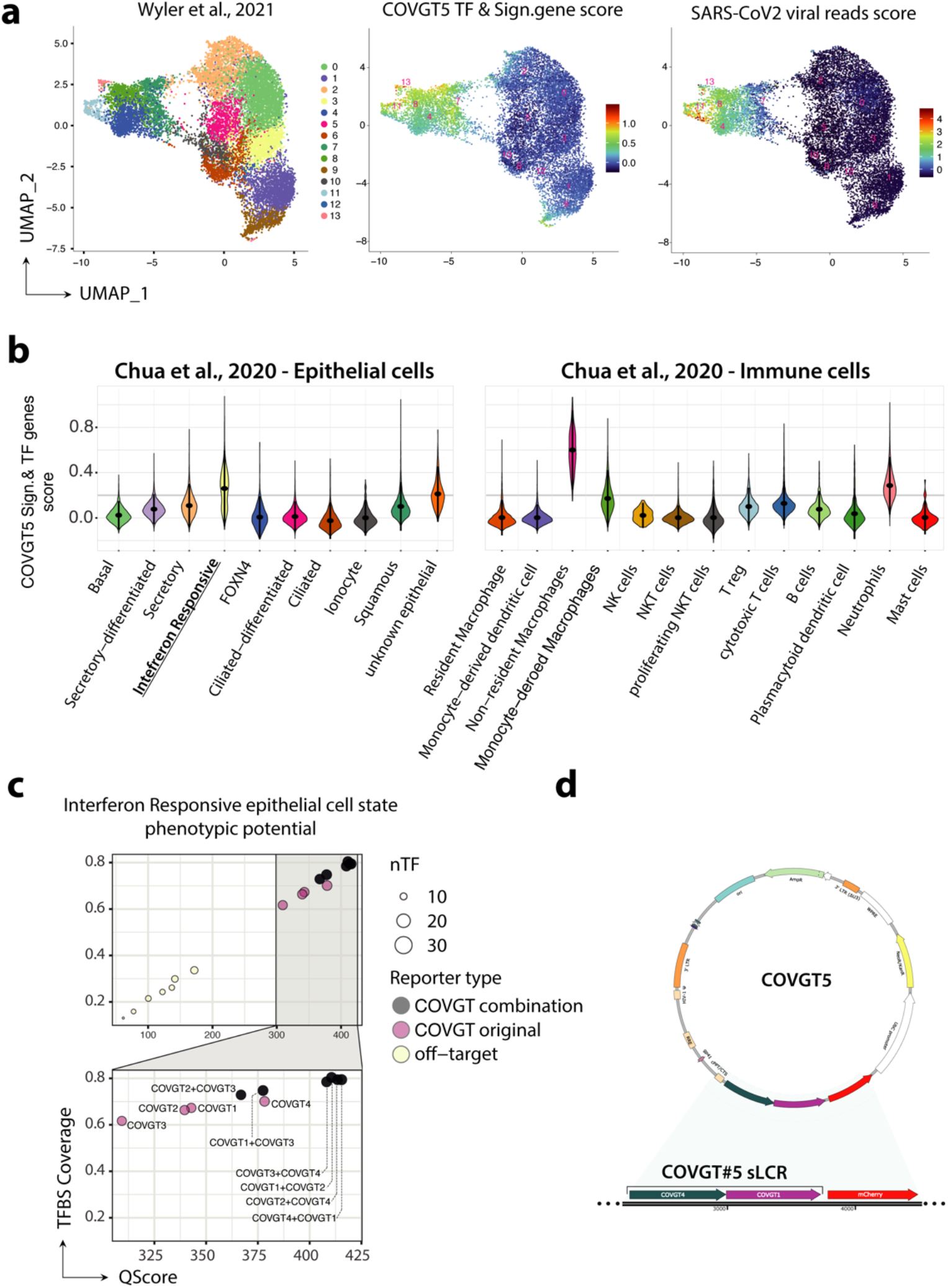
COVGT5 represents a productive SARS-CoV-2 infection and innate immune response *in silico*. scRNA-seq clustering of SARS-CoV-2. UMAP dimensional reduction of data from Wyler et al. 2020 SARS-CoV/CoV-2. Colors highlight cell clusters enriched for SARS-CoV2 viral reads and COVGT5 signature scores. **(B)** Violin plot ranking COVGT5 against SARS-CoV-2 patient samples. Signature and transcription factor gene scores from COVGT5 were calculated on annotated cell clusters from Chua et al. 2020. Enrichment scores (y-axis) of distinct cell clusters (x-axis) within epithelial (left) and immune (right) cell populations are shown. **(C)** *In silico* comparison of individual and combined COVGT sLCRs. Six combinations of sLCRs (black) are plotted against individual COVGT sLCRs (pink) and off-target reporters (yellow). Qscore (x-axis) and TBFS coverage (y-axis) predict specificity on the target phenotype. The dot size indicates the number of transcription factors captured from the input list. **(D)** Vector composition of COVGT5-mCherry. Representation of COVGT5 design from a combined COVGT4-COVGT1 sLCR driving mCherry expression.

To advance reporters that may be representative of primary cell responses in patients, we ranked all the reporters against the nasopharyngeal and bronchial samples from 19 clinically well-characterized SARS-CoV-2 patients with moderate or critical COVID-19 and five healthy controls ^7^. Interestingly, the SARS-CoV-2 signature genes retrieved in three independent cellular models for SARS-CoV-2 infection and underlying the design of two distinct sLCRs (COVGT1 and COVGT4) identified two primary cell populations: myeloid cells, notably inflammatory macrophages, and one subpopulation of ciliated epithelial cells with a distinctively strong interferon gamma (IFNG) response signature (**Fig. 2b** and **Fig. S2a-b**). This is consistent with the pathway analysis of the signature and TF genes (**Fig. 1b-c**) and suggested that COVGT1 and COVGT4 may cumulatively represent host primary cells in which viral infection triggered an innate immune response. Since COVGT1 and COVGT4 individually display a mild but specific response to IFN stimulation and combinatorically marked a population of epithelial cells activated by SARS-CoV-2 in patients (**Fig. 2c**). To aim at the highest activity potential, we combined these two reporters into one, hereafter referred to as COVGT5 (**Fig. 2d**).

### COVGT5 responds to SARS-CoV-2 and triggers of epithelial innate immunity

To validate the designed reporters, we genetically engineered using transposon-based vectors Calu-3, A549 and 293T cells, which represent a limited but diverse set of lung cancer and immortalized kidney cells lines. We next tested by FACS and immunofluorescence the individual expression COVGT1-5 sLCRs and found that COVGT5 readily drove the fluorescent protein expression in response to a mix of interferons of all classes (hereon referred to IFN mix; **Fig. 3a**). Such response was markedly more pronounced in non-transformed kidney cells than in lung cancer lines. Calu-3 responsiveness to interferons was especially weak (**Fig.3a** and **s3a-c**). COVGT1-4 sLCRs typically had low basal expression that is compatible with screening for inducers (**Fig. 3a-c, s3a-c** and methods) and low inducibility (**Fig. S3a**), consistent with their phenotypic potential being narrower than COVGT#5 (**Fig. 2c**).

**Fig. 3.**
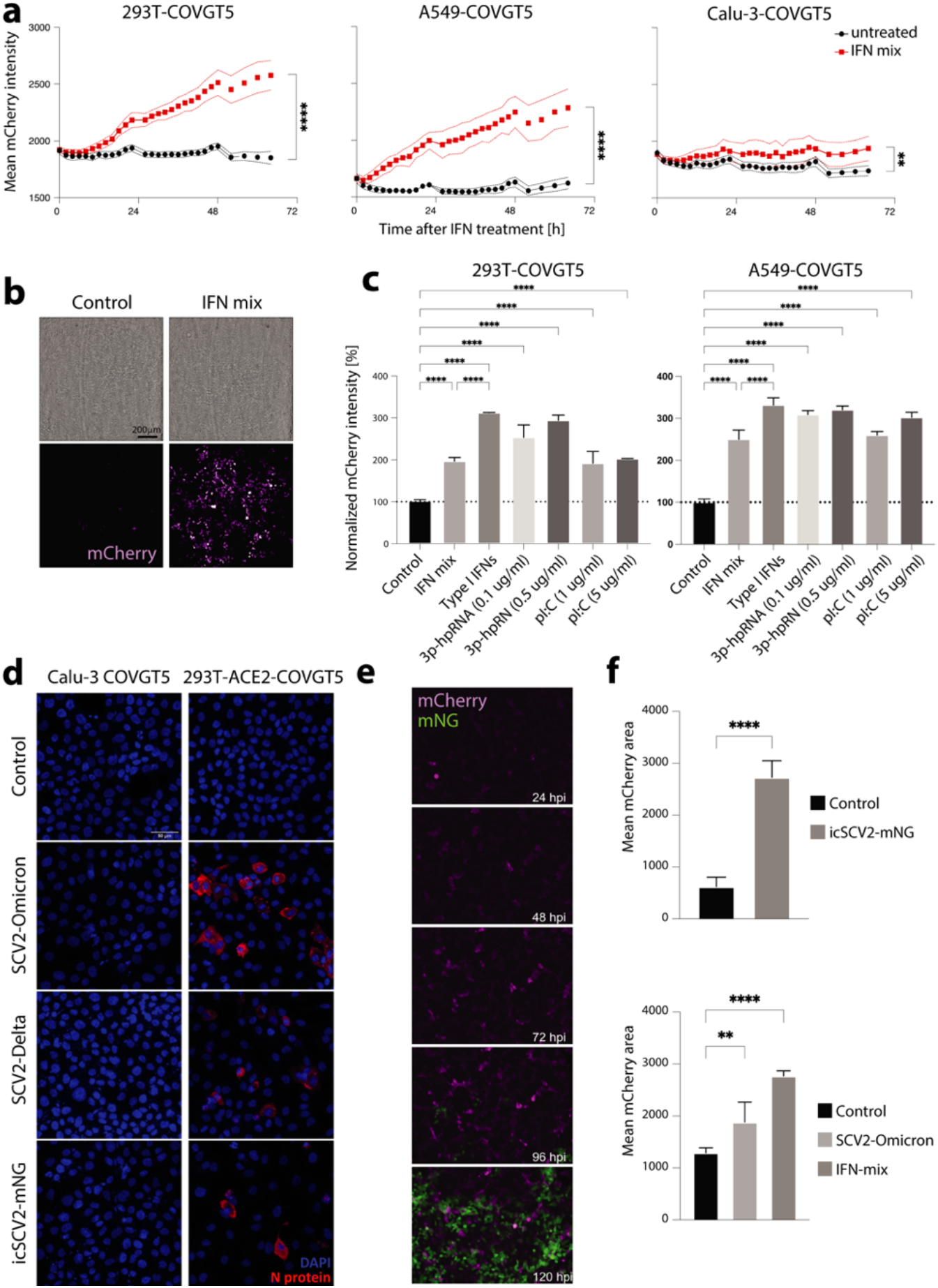
COVGT5 responds to SARS-CoV-2 and innate immunity activators. **(A)**Longitudinal measurement of 293T, A549, and Calu-3 COVGT5 reporter cells upon IFN mix treatment. Time after treatment is indicated on the x-axis, each dot represents a single measurement. Mean mCherry intensity was calculated from technical triplicates (n=3) measured by Operetta confocal imaging system and the confidence interval was plotted. **(B)** Representative images of COVGT5-driven mCherry expression (purple) in 293T. Images were taken 48h after IFN mix treatment. The scale bar indicates 200µm. **(C)** Bar plot quantification of COVGT5-driven mCherry upregulation. COVGT5 induction at 48h following IFN mix, 3p-hpRNA (0.1ug/ml, 0.5ug/ml) and pI:C (1ug/ml, 5ug/ml) treatment in 293T and A549 reporter cells was assessed by normalizing against untreated reporter cells. P-values denote significance by 1-way ANOVA. **(D)** Fluorescence microscopy images of SCV2 infection. N-protein (red) was used as a proxy for productive SCV2 infection with Omicron, Delta, and mNG-engineered Wuhan strain in Calu-3 and 293T-ACE2 reporter cells. Nuclei are counterstained with DAPI (blue). **(E)** Longitudinal fluorescence microscopy images of COVGT5-driven mCherry expression (purple). 6600pfu/ml icSCV2-mNG (green) were used to infect 293T-COVGT to visualize productive infection. **(F)** Bar plot quantification of COVGT5-driven mCherry upregulation in response to 6600pfu/ml icSCV2-mNG and 88pfu/ml SCV2-Omicron. Measurement by Incucyte were at 120h post-infection. The relative mean fluorescent area was determined from technical replicates (n=4). P-values denote significance by 1-way ANOVA.

sLCRs may be used to discover external signaling leading to phenotypic state transitions ^14,15^. Thus, we next tested specific cues that may potentially drive the response of COVGT5 and be relevant for viral infection biology. Importantly, COVGT5-modified 293T and A549 cell lines readily responded to individual stimulation by type I interferons, IFN-α-2a, IFN-α-2b and IFN-β, as well as synthetic double-stranded RNA transfection but not to IFN-g, IL-28 (IFN-Α), TNFa or LPS (**Fig. 3c and s3d**). The marked response to short, tri-phosphorylated stem-loop hairpin RNAs (3p-hpRNA) as compared to the less specific inducer of PRRs, polyinosinic/polycytidylic acid (poly(I:C)), indicated that RIG-I-mediated innate intracellular double-stranded RNA sensing ^20^ is a key trigger of COVGT5 expression.

Next, we tested COVGT5 induction in response to SARS-CoV-2 infection. Whereas Calu-3 cells endogenously express the SARS-CoV-2 human receptor ACE2 and the TMPSSR2 protease, albeit at low levels, the other cell lines are predictably resistant to infection. Therefore, we engineered A549-COVGT5 and 293T-COVGT5 lines to over-express human ACE2. Extracellular ACE2 enzymatic activity confirmed the successful engineering (**Fig. S3e**) and made both cell lines permissive for infection with all the strains tested (Wuhan, Delta and Omicron; **Fig. 3d**). Furthermore, introduction of ACE2 did not affect COVGT5 reporter inducibility in 293T and A549 exposed to the IFN mix (**Fig. S3f**). In a longitudinal live-cell imaging setting using the wild-type Wuhan SARS-CoV-2 strain engineered to express mNeonGreen (icSCV2-mNG ^21^), A549-ACE2-COVGT5 and 293T-ACE2-COVGT5 lines were permissive to infection and a measurable viral accumulation/replication starting from 72-120 hours post-infection (hpi; **Fig. 3e**). COVGT5 response appeared to be proportional to the extent of viral replication in our cell models. A high dose of icSCV2 caused a marked response in 293T-ACE2-COVGT5, which was comparable to interferon stimulation (**Fig. 3f**). Consistently, in response to low-dose Omicron infection, we observed a moderate but measurable activation of the COVGT5 sLCR, which was delayed and reduced when compared to their response to interferons (**Fig. 3f and s3g**).

In our experiments, Calu-3 cells, despite being initially adopted by the field as model of choice for *in vitro* infection biology, proved to be less responsive to interferon signaling (**Fig. 3a and s3b-c**) and appeared poorly permissive for infection with all the strains tested, which was confirmed by SARS-CoV-2 N-protein immunofluorescence (**Fig. 3d**) Overall, our experiments show that COVGT5 responds to type I interferons, double-stranded RNA, and SARS-CoV-2 viral replication, which are all triggers of epithelial innate immunity.

### A platform for discovery of pharmacological modulators of cellular innate immunity and response to SARS-CoV-2 infection

To exploit the specific responses of COVGT5 to triggers of epithelial innate immunity, we next subjected our epithelial cell models to the above-validated signaling cues and systematically investigated pharmacological modulators of such response.

To this end, we assembled a custom library of small molecules against some of the key targets and pathways, including targeted agents approved by the United States Food and Drug Administration (FDA; Supplementary Table S2). To screen for modulators of epithelial cell response to COVGT5 activation that may be used in clinics, we used a treatment scheme including three doses in the low-nanomolar to -micromolar range (i.e. 0.03, 0.3, 3 mM; **Fig. 4a-b and s4a**). In a forty-eight-hour live imaging setting, we were able to identify drugs that would affect cell viability even at low doses/short timepoints and rank selected drugs for their ability to enhance or block COVGT5 activation in response to both IFN mix and 3p-hpRNA. Importantly, the top ranked small molecules that blocked COVGT5 activation in both lines and in response to both triggers, were well established and clinically relevant JAK inhibitors Baricitinib and Tofacitinib (**Fig. 4b-c and s4b**). The mechanism of action of both drugs is the inhibition of autocrine and paracrine cytokine signaling, with specificity for JAK1-JAK2 (Baricitinib) and JAK3-JAK2 (Tofacitinib), in agreement with the JAK-mediated STAT1 activation caused by type-I IFN stimulation of the IFNAR. Since COVGT5 activation was efficiently blocked by both JAK inhibitors in both cell lines in response to the JAK1-/STAT1-independent trigger 3p-hpRNA, the screen suggests that COVGT5 reports on signaling of both innate and adaptive immune responses in epithelial cells.

**Fig. 4.**
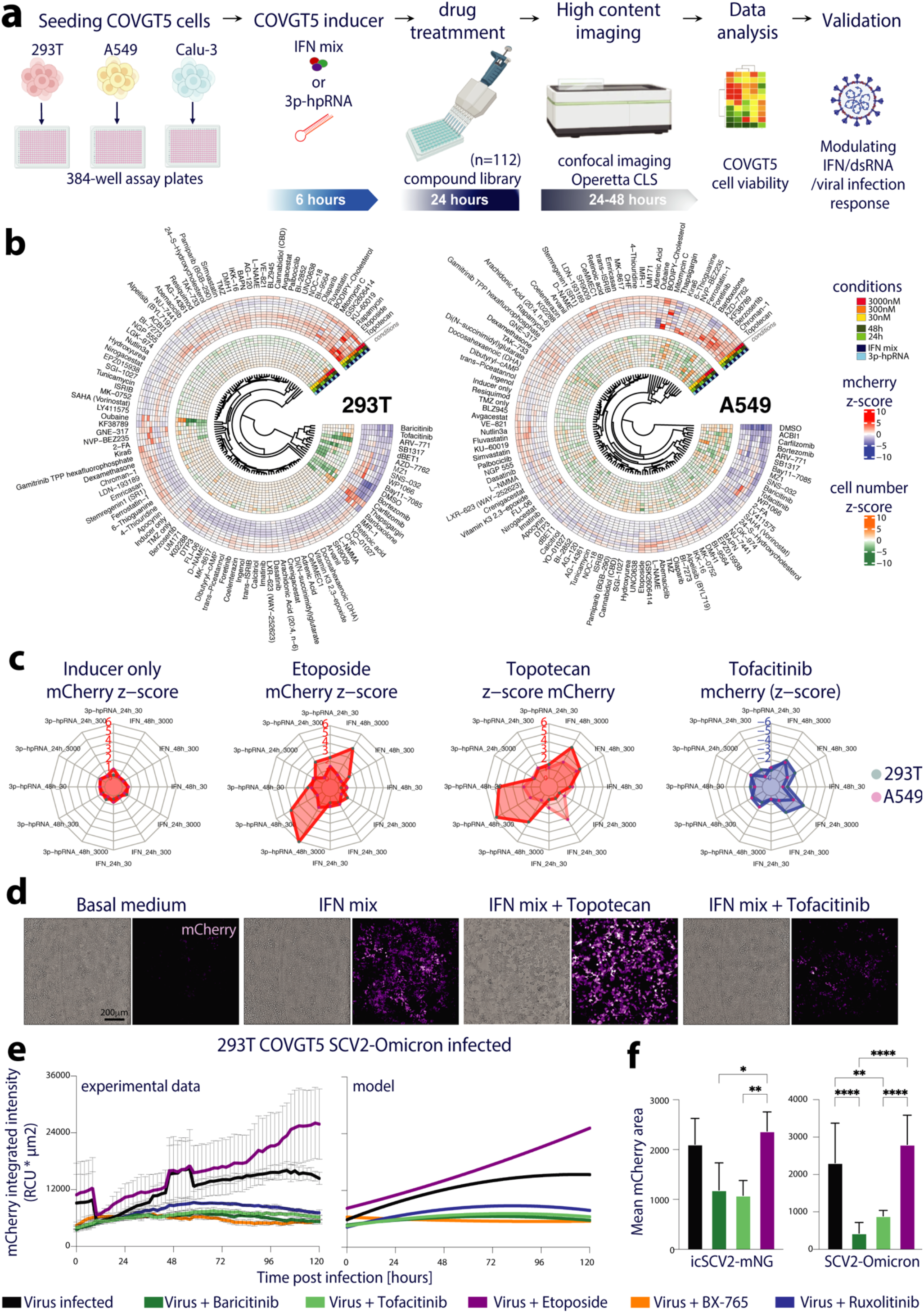
A platform for discovery of pharmacological modulators of cellular innate immunity and response to SARS-CoV-2 infection. **(A)** Schematic representation of the platform. 112 pharmacological agents were tested on 293T, A549, and Calu-3 COVGT5 reporter cells using IFN mix and 0.1ug/ml 3p-hpRNA as inducers and Operetta image acquisition and quantification as readout. **(B)** Circular heatmaps representing screening results of 293T (left) and A549 (right) reporter cells. Drugs were tested in concentrations 30nM, 300nM, and 3000nM at 24h and 48h post-induction in technical replicates (n=4). Z-scores calculated for cell viability (inner circle) and mCherry intensity (outer circle) are plotted and drugs are ordered based on hierarchical clustering for both parameters. **(C)** Spider plots highlighting COVGT5 modulators. Amplifiers (Topotecan and Etoposide) and inhibitors (Tofacitinib) were identified from screening data in 293T (teal) and A549 (purple). Each data point is represented as a corner of the plot with the indicated scale ranges (note color coding for red=activation and blue=inhibition). **(D)** Representative images showing modulatory drug effect. 300nM Topotecan and 300nM Tofacitinib were used on top of the IFN mix in 293T reporter cells. The scale indicates 200um. **(E)** Longitudinal measurement (left) of COVGT5 modulation in virus infection setting. 293T-COVGT5 were infected with 88 pfu/ml SCV2-Omicron and treated with 10uM Baricitinib, 10uM Tofacitinib, 10uM Etoposide, 200nM BX-765 or 2uM Ruxolitinib. Polynomial regression model (right) of longitudinal measurement pronouncing drug effects on COVGT5 induction. Measurement and quantification were performed using Incucyte factoring mean fluorescence intensity and fluorescent area. **(F)** Independent bar plot quantifications of 293T-COVGT5 induction by SCV2 infection. 6600pfu/ml SCV2-Omicron or icSCV2-mNG were used with Baricitinib, Tofacitinib, and Etoposide at 120h post-infection. Measurement and quantification were performed using Incucyte. The relative mean fluorescent area was determined from technical replicates (n=4). P-values denote significance by 1-way ANOVA.

Strikingly, among the top amplifiers of COVGT5 activation in response to the aforementioned triggers and across multiple cell lines and doses, we scored several modulators of genome integrity, with the notable case of DNA damage inducers (e.g. TOP1 inhibitor, Topotecan and TOP2 inhibitor, Etoposide; **Fig. 4b-c and s4b**). The main hits of the screens were independently validated in the reporter cell lines, and specific COVGT5 expression was confirmed (**Fig. 4d**).

### Etoposide and Tofacitinib antagonistically modulate COVGT5 response to productive SARS-CoV-2 infection

To validate the response observed in the screen using SARS-CoV-2, we next infected 293T-ACE2-COVGT5 with the Omicron strain and simultaneously treated these cells with inhibitors of either TOP2 (Etoposide), JAK (Baricitinib, Tofacitinib and Ruxolitinib), or TBK-1/IKK-ε/PDK-1 (BX-795). The latter blocks noncanonical IκB kinase and IRF-3 activation during innate immune sensing. During the longitudinal monitoring of cell responses, 293T-ACE2-COVGT5 displayed a measurable activation of the COVGT5 sLCR in response to Omicron infection, with a moderate upward trajectory at the pre-defined time point of 156 hpi. This activation mode appeared to be specific for the viral infection setting, as cellular response to the interferon signaling reached the plateau at 48 hours post cytokine stimulation and lasted until 156 hpi (**Fig. S4c**). Importantly, in line with the results of our screen, COVGT5 expression was completely abrogated by all the independent JAK inhibitors tested (Ruxolitinib, Baricitinib and Tofacitinib), as well as by the TBK-1/IKK-ε/PDK-1 inhibitor BX-795 (**Fig. 4e**). Instead, Etoposide imparted a moderate amplification of the COVIGT#5 signal when administered in parallel to Omicron (**Fig. 4e**), and a nonlinear regression model supported the hypothesis that Etoposide and JAKi antagonistically modulate COVGT5 response to productive SARS-CoV-2 infection (**Fig. 4e**). Similar antagonism was observed for the SCV2-mNG strain (**Fig. 4f and s4c**).

Thus, our data support the use of a sLCR to detect responses to viral infection as a drug discovery platform to identify pharmacological modulators of innate immune responses.

### Synergistic and antagonistic pharmacological modulation of COVGT5 underscored drug mechanism of action

The targets of individual JAK inhibitors are biochemically well established and the consequences of signal transduction observed in our experiments aligned with expectations. Cell exposure to chemotherapeutics such as Topotecan and Etoposide leads to far less predictable targets and consequences. We therefore tested if our reporter could provide evidence supporting the mechanism of action of the drugs controlling its regulation.

DNA damage induces IFN-stimulated genes and the IFN-α and IFN-Α genes via NF-kB activation ^12^, which is consistent with both the predicted outcome of TOP1 and TOP2 inhibition by Topotecan and Etoposide, respectively. However, small molecules may also have unrelated, off-target effects. To address if DNA damage is the actual trigger of the SARS-CoV-2 infection-like response observed, we set out to conduct selective pairwise synergy screens. In 293T-COVGT5 and A549-COVGT5 exposed to IFN mix or 3p-hpRNA, Bliss synergy scoring of the dose-escalation for Topotecan and Etoposide determined that the combined treatment synergistically amplified COVGT5 expression, in line with their predicted induction of DNA damage (**Fig. 5a** and **s5a**). The extent of synergism was cell line specific and reflected the extent of response to IFN mix (**Fig. 5a** and **s5a**). Conversely, JAK inhibitors combination synergistically restricted COVGT5 activation (**Fig. 5a** and **s5a**), indicating that synergism reveals mechanistical convergence. Thus, we next substituted either Topotecan or Etoposide with ionizing radiation (IR), which is a direct method to damage DNA. IR amplified activation of COVGT5 in dose-dependent manner on top of 3p-hpRNA transfection, and in both lung and kidney cell lines, starting from a dose of 1 gray (1 Gy; **Fig. S5b**). Importantly, Bliss synergy scoring of the dose-escalation of each combination supported that IR activation is epistatic to Topotecan or Etoposide (**Fig. 5b**), supporting that cooperative DNA damage induction occurred. In contrast, Bardoxolone, an NRF2 activator, antagonistically regulated COVGT5 when combined with Topotecan (**Fig. S3c**), indicating that drugs with distinct mechanisms of action do not synergistically amplify cellular responses to 3p-hpRNA.

**Fig. 5.**
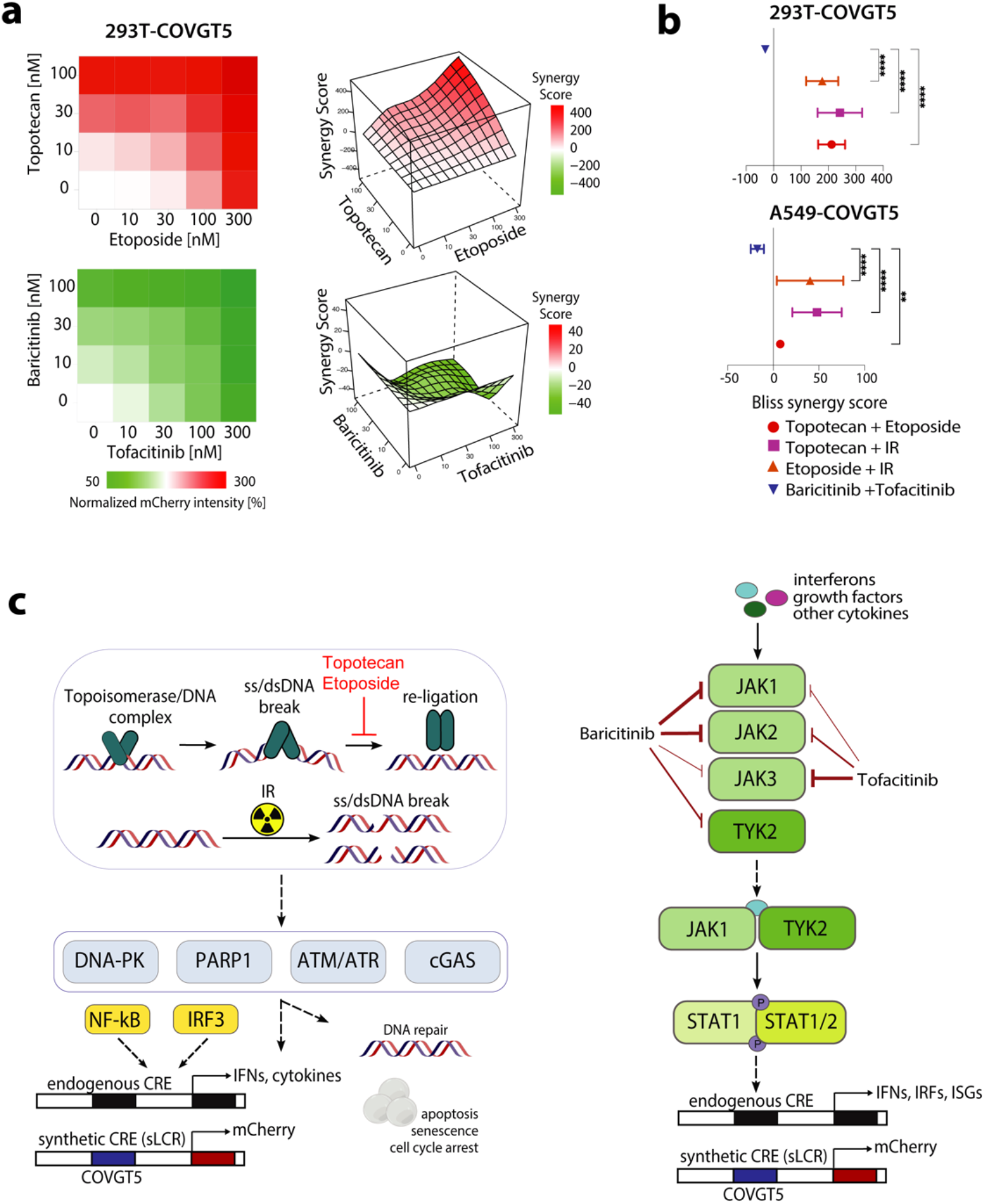
Synergistic and antagonistic pharmacological modulation of COVGT5 underscored the drug mechanism of action. **(A)**Dose-escalation of Topotecan against Etoposide (top) and Baricitinib against Tofacitinib (bottom). 293T-COVGT5 received IFN mix induction followed by combinatorial drug treatment. Mean mCherry intensity was calculated from technical triplicates (n=3) and normalized to non-drug treated cells. Quantification was performed by high throughput FACS measuring mean mCherry fluorescence intensity. Bliss synergy scores inform on synergism between Topotecan + Etoposide and Baricitinib + Tofacitinib in modulating COVGT5 response. **(B)** Mean Bliss synergy score calculated from all single drug combination data points. Topotecan + IR and Etoposide + IR in addition to drug combinations from a) in 293T and A549 reporter cells were performed in technical triplicates (n=3). Quantification was performed by high throughput FACS. P-values denote significance by 1-way ANOVA. **(C)** Schematic representation of proposed mechanisms of action for COVGT5 modulation. Pathways for induction and amplification by Topotecan, Etoposide, and IR (left) and COVGT5 inhibition by Baricitinib and Tofacitinib (right) (strength of inhibition arrows indicate specificity) are proposed.

Altogether, our data suggest that DNA damaging agents and JAKi antagonistically modulate epithelial innate immunity in response to SARS-CoV-2 infection (**Fig. 5c**). These leads were enabled by synthetic genetic tracing for SARS-CoV-2 viral infection in epithelial cells applied in a drug discovery setting and feature its application to infection biology more broadly.

## Discussion

COVID-19 pandemic has seen no single drug or combination proven to effectively halt disease progression, and the effect of the massive worldwide vaccination campaign is waning. Near future surges of this pandemic by Omicron and future subvariants, as well as pandemic-scale infectious diseases are recurrent in history. Here, we report the COVGT sLCRs, and the validation of the approach to design such reporters, as a useful resource for investigating mechanisms of cellular response to SARS-CoV-2 entry and replication as well as a phenotypic platform for high-throughput screening of therapeutics. Efforts to combine reporters for viral entry and replication with those measuring cellular transcriptional responses are timely ^22^. The flexibility to use such systems in any chosen model, potentially including preclinical models *in vivo*, is a unique feature of our system. Additionally, the adaptability of our system to establish *in vitro* infection models allows the rapid testing of wild-type and evolving pathogens stains. Whereas heterogeneous responses to viral infection are likely rooted in biology ^22^, future experiments with this and other reporters will likely benefit from using cellular models that are physiologically more accurate than cell lines. Incidentally, our experiments uncovered 293T cells as a better responsive model to triggers of innate immunity and viral infection. We generally observed intermediate interferon and RIG-I responses as well as SARS-CoV-2 infection permissiveness in A549, and poor responses in Calu-3, a lung cancer cell line quickly adopted by the field during the early phases of the pandemic response, and later dismissed.

The herein characterized COVGT5 reporter outperformed all broadly available reporters *in silico* and is independently and strongly activated by type I interferons and RIG-I agonists in cells. RIG-I, as a major receptor for viral mRNA sensing, was validated head-to-head in primary cells and cell lines ^23^, supporting the validity of our findings. Such activation in our epithelial cell models may be modulated by bioactive agents discovered through *in vitro* image-based screens, as exemplified by our discovery of DNA damage inducers and JAK inhibitors as rheostats of epithelial immunity. Of note, the in silico single-cell analysis suggests that COVGT5 responses may be conserved across tissues

Interferon response constitutes one of the first lines of defense against viral infections. It involves the transcription of interferon genes first and later of interferon-stimulated genes (ISGs), which in turn lead to an antiviral state in both infected and neighboring cells. Viral infections are detected through pattern recognition receptors, including RIG-I-like receptors. Importantly, SARS-CoV-2 evolution through variants of concern involved increased interferon response evasion ^24^ and a synthetic agonist stimulation of RIG-I protects mice from acute and chronic SARS-CoV-2 infection, even in absence of the adaptive immune system ^25^. This underscores the importance of epithelial innate immune responses to restrain SARS-CoV-2 infection and supports the identification of pharmacological modulators of such response as a therapeutic strategy for COVID-19 and viral infection more broadly. Our COVGT5 sLCR designed on the transcriptional response of lung epithelial and enterocyte organoids to SARS-CoV-2 infection could be explained by activation of interferon-and RIG-I-like signaling. This is an additional asset for our study that provides a cellular reporter for viral infection biology and a molecular tool for cellular and molecular dissection of these pathways.

Given the antiviral immunity’s pivotal signaling through JAK/STAT pathway, JAK inhibitors were already clinically tested as therapeutics for COVID-19. Retrospectively, JAKi appeared potentially helpful to manage disease severity ^3^. Likewise, Topotecan, one of the other drugs uncovered by our COVGT5 reporter as an amplifier of innate immunity, was recently discovered as treatment to blunt the hyper-inflammatory response to anti-SARS-CoV-2 treatment with successful validation in preclinical models^26^. More generally, Topoisomerase 1 and 2 inhibitors are potent modulators of innate immune activation ^27,28^. An ongoing clinical trial is testing Topotecan in COVID-19 patients to assess the safety of a single dose and its potential to blunt inflammation systemically (NCT05083000). Our results may be helpful to interpret the results of this trial as they suggest that DNA damage modulation may also be effective in priming an antiviral state in vivo, or amplifying it. This may occur in both infected and neighboring cells as a function of cellular state or productiveness of the infection.

Combination therapies were frequently tested in COVID-19 trials, in particular using antiviral drugs as anchor, since those alone were deemed insufficient if administered at disease progression ^5^. Remdesivir, the first antiviral considered mildly effective against COVID-19, appeared to cooperate with Baricitinib ^5^. For both JAK inhibitors and dexamethasone, a glucocorticoid receptor agonist that did not constantly modulate our reporter across screens and their validations, trials returned the cautionary note that the drug administration should occur in a particular time window promoting their beneficial effects, or could otherwise become detrimental ^3,4^. In our experiments, all JAK inhibitors abrogated cellular response to viral replication supporting their use to that effect. We arrived to conclusions through rapid screening of a curated set of drugs, which otherwise required the combination of artificial intelligence and clinical trials{Stebbing:2021db}. Moreover, our platform is uniquely placed to discover drug combinations synergistically acting toward cellular response to viral infection in preclinical models, which may later be tested during disease progression. Alternatively, rather than abrogating epithelial immunity, drugs synergistically promoting an antiviral state that cooperates with antiviral drugs could be tested to potentiate disease control. As supplementing interferon beta-1a in combination with Remdesivir did not improve outcomes and was associated with adverse effects in a placebo-controlled phase III trial ^29^, innate immunity amplification will require the identification of safe combinations. Our screen also initially revealed Rapamycin as a potential epithelial immunity amplifier drug. However, this drug was not validated in later experiments. Whether targeted ionizing radiation may reduce the impact of viral shedding in selected organs is an intriguing hypothesis that stems from this study. If DNA damage-inducing agents in patients with COVID-19 and cancer may be consider feasible, their potential side effect of inducing senescence thereby exacerbating cellular responses to infection has to be ruled out preclinically ^30^. Moreover, future drug screening combinations may focus more systematically on approved drugs with higher safety profiles. Yet, therapies amplifying interferon response should involve *in vivo* testing in preclinical models and consider that this pathway is implicated in severe COVID-related complications, such as vasculitis, “COVID toes” and Kawasaki-like disease. Regardless, our reporter will be a valuable tool to mechanistically dissect the endogenous gene regulatory networks of innate immunity in epithelial cells and beyond.

The onset of the COVID-19 pandemic mounted into substantial global morbidity and mortality, and worldwide economic collapse, due to the lack of early preparedness. Our approach required four weeks from availability of the first transcriptional signatures to reporter validation and holds outstanding potential for optimization, thereby supporting synthetic genetic tracing for infection biology more readily in the future.

## Materials and Methods Datasets

The data to generate the signature genes and transcription factors lists in Fig.1 were downloaded from Gene Expression Omnibus, accession GSE148729 and GSE147507, or supplementary material (see below).

### LSD reporters design

COVGT reporters were designed using the LSD method, as described in (Company C., Schmitt M.J., Dramaretska Y., et al., *manuscript in preparation*).

Briefly, the LSD algorithm takes a list of PWMs, a list of marker genes of a target phenotype, and the reference genome of the organism of interest, and it generates a list of naturally-occurring, putative *cis*-regulatory elements used to assemble the synthetic-reporter. The algorithm can be divided into three steps. In step I, LSD generates a pool of potential CRE with a fixed length within user-defined regulatory landscapes (default is a 150bp window sliding with a 50bp step). In step II, LSD assigns TF-binding sites to the CRE pool using FIMO (default *--output-pthresh 1e-4 --no-qvalue*), and creates a matrix of putative CREs x TFBS. In step III, LSD ranks and selects the minimal number of CREs representing the complete set of TFBS. For that purpose, it uses an algorithm to sort and select the best CRE based on the overall TFBS affinity and diversity among input TFs showing high affinity for the CRE. Starting from the ranked CREs, LSD selects the highest-ranking CRE defined by the sum of the affinity score (-log10(p-value)) and TFBS diversity (number of different TFBS). Subsequently, it removes the selected CRE and the corresponding TFBS from the CRE x TFBS matrix and repeats the selection. This continues until either none of the CRE or of the TFBS is left. In the ranking, priority is given to CREs proximal to known TSS, based on 5’ CAGE data (ENCODE) in order to increase the chances of successful transcriptional firing using the same strategy as above. Finally, LSD returns an ordered list of the selected CREs, together with a representation of the TFBS scores (**Figure 1A**). LSD is available at: https://gitlab.com/gargiulo_lab/sLCR_selection_framework.

### Cis-regulatory potential assessment of the SARS-CoV-2 sLCRs

In order to generate a common SARS-CoV-2 signature and define the specific TF (**Fig 1a**), we integrated different transcriptional profiles available at the moment of this publication. We first downloaded Blanco et al. 2020 (GSE147507), Wyler et al. 2020 (GSE148729; www.mdc-berlin.de/singlecell-SARSCoV2), and Lamer et al. 2020 (Differentially expressed tables). Then, we reanalyzed Blanco et al. 2020 and Wyler et al. 2020 using the same parameters as in the publications. Blanco et al. 2020 and Lamers et. al. 2020, gene signatures and TF were selected using log2FC > 1 and adj.p-value < 0.05. Wyler et al. 2020 gene signatures were identified by using the FindMarkers function (Seurat v.3.1, R v.3.6), and selected using < 0.05 adj.p-value as a threshold. scRNA-seq TF selection was generated by selecting exclusive highly-expressed (quantile >75%) TF genes. Integration of the data-sets to construct the common gene signatures was done using at least three common genes between experiments, and two common TF genes to define the TF input. Further analysis of TF using Ingenuity pathway analysis (IPA) generated the final set of TFs and TFBS used in the different comparisons (**Fig. S1**). Previously published reporters carrying promoters of the human interferon gamma or beta genes were included as benchmark (Addgene #102597, #17596, #30536 and #17598). SARS-CoV-2 sLCRs pool was designed using a combination of individual and common data-sets and the LSD framework. The inference of the phenotypic potential using was generated as described above, but using as reference IPA selected TFBS. Heatmaps and gene set enrichment analysis (**Fig. 1b-c)** were generated using common genes and selection with pheatmap v.1.0.12 and piano v.2.0.2, respectively. Hg19 assembly was used to extract references and generate the sLCR vectors. Scatter plots graphics were generated using ggplot2 on an R v.3.6 environment.

### Signature scoring on single-cell RNAseq datasets

Two single-cell RNAseq studies were used to establish scores for the COVGT5 signature list consisting of marker genes and transcription factors. From Wyler et al. (16), we downloaded the Seurat object file and retrieved annotations for viral reads (SCoV1-ORF1a, SCoV1-ORF1ab, SCoV1-S, SCoV1-ORF3a, SCoV1-ORF3b, SCoV1-E, SCoV1-M, SCoV1-ORF6, SCoV1-ORF7a, SCoV1-ORF7b, SCoV1-ORF8a, SCoV1-ORF8b, SCoV1-N, SCoV1-ORF10, SCoV1-UTR3, SCoV2-orf1ab, SCoV2-ORF10, SCoV2-ORF3a, SCoV2-E, SCoV2-M, SCoV2-ORF7a, SCoV2-ORF8, SCoV2-N, SCoV2-UTR3. Accession numbers: AY310120 for SARS-CoV, MN908947 for SARS-CoV-2). The list of viral reads as well as the list of signature genes and transcription factors for COVGT5 generation were used to calculate the meta-module score on single cells using the AddModuleScore() function from Seurat. Similarly, from Chua et al. (7) and https://digital.bihealth.org, we downloaded the Seurat object and used the AddModuleScore() function to determine enrichment of the COVGT5 signature list over the annotated cell clusters.

### Cell lines

All lines used in this study were thawed from frozen batches and propagated for a limited number of passages (10-15×), and all lines regularly tested with the Mycoplasma Detection Kit (Jena Biosci-ence; 11828383, PP-401L) to exclude contamination. 293T and A549 cell lines (R. Bernards laboratory, Netherlands Cancer Institute (NKI), Amsterdam, Netherlands) were cultured in DMEM or RPMI medium (Gibco) respectively, supplemented with 10% fetal bovine serum (FBS, from Gibco) and penicillin and streptomycin (100 U/ml; Gibco) at 37°C in a 5% CO_2_ and 95% air. Calu-3 (L. Čičin-Šain laboratory, Helmholtz Centre for Infection Research (HZI), Braunschweig, Germany) were cultured in DMEM medium (Gibco), supplemented with 1x MEM NEAA (Gibco), 1x sodium pyruvate (Gibco), 10% fetal bovine serum (FBS, from Gibco) and penicillin and streptomycin (100 U/ml; Gibco) at 37°C in a 5% CO_2_ and 95% air.

### Transfection/Transduction

Transfection and transduction were previously described in detail (11). Briefly, 12μg of DNA mix (lentivector, pCMV-G, pRSV-REV, pMDLG/pRRE) was incubated with the FuGENE (Promega, E2311)– DMEM/F12 (Life Technologies, 31331) mix for 15 minutes at room temperature, added to the antibiotic-free medium covering the 293T cells, and the first tap of viral supernatant was collected at 40 hours after transfection. Titer was assessed using the Lenti-X p24 Rapid Titer Kit (Takara, 631280) according to the manufacturer’s instructions. We applied viral particles to target cells in the appropriate complete medium supplemented with 2.5μg/mL protamine sulfate. After 12 to 14 hours of incubation with the viral supernatant, the medium was refreshed with the appropriate complete medium.

### COVGT reporter cell lines

COVGT1-4 reporter cell lines were generated by PiggyBac vector delivery: 3µg of Super PiggyBac Transposase (System Biosciences, PB210PA-1-SBI) and 3µg of reporter plasmid (pPB-{COVGT1}>d2EGFP, pPB[-{COVGT2}>d2EGFP, pPB-{COVGT3}>d2EGFP, pPB-{COVGT4 }>d2EGFP) were transfected. Transfected cells were selected using 1mg/ml G418 (Sigma). COVGT5 reporter cell lines were generated through lentiviral transduction with pBA407-{COVGT5}>mCherry. Infected cells were selected using 1mg/ml G418 (Sigma). ACE2 expression 293T and A549 reporter cell lines were generated through lentiviral transduction with pscALPSpuro-HsACE2 (human) (Addgene, #158081). Infected cells were selected using 2µg/ml puromycin (Sigma).

### FACS sorting

Transduced cell lines were harvested into single-cell suspensions and resuspended into cold medium and filtered through 40µm mesh filters (BD) into FACS tubes. Sorting was conducted using BD FACSAria III or Fusion. The appropriate laser-filter combinations were chosen depending on the fluorophores being sorted for (GFP or mCherry). Typically, to remove dead cells, events were first gated on the basis of shape and granularity (FSC-A vs. SSC-A) and doublets were excluded (FSC-A vs. FSC-H). If insufficient, alternative gates were used (FSC-W and SSC-W). Positive gates were established on parental cells, untreated reporter cells and IFN mix induced reporter cells, to sort for populations with fluorescent signal above background (Fig. S3a-b, f).

### COVGT reporter induction

For COVGT reporter induction, cells were seeded at a density of 2000 in 50µl (for 384-well plates, Greiner) or 10,000 in 100µl (96-well plates, Greiner) cells per well and incubated at 37°C in a 5% CO_2_ and 95% air for 1-2 days until a confluency of 60-80% was reached. Cells were treated with individual cytokines (Peprotech) or COVGT induction mixes: Inflammation medium – TNFa (5ng/ml), IFNg (10ng/ml), IL4 (10ng/ml), EGF (10ng/ml), IL6 (10ng/ml). IFN mix – IFNa-2a (20ng/ml), IFNa-2b (20ng/ml), IFNb-2b (20ng/ml), IL-28 (IFNΑ; 10ng/ml), IFNg (5ng/ml). 3p-hpRNA, pI:C (Invivogen) and LPS (Sigma) – Transfected at indicated concentrations following transfection protocol described above. Quantification of reporter expression was performed by FACS or Operetta CLS typically 48h following induction.

### Fluorescence microscopy

Operetta CLS (Perkin Elmer) and Incucyte (Sartorius) were used to acquire fluorescence imaging data of COVGT5 induction. Reporter cell lines were plated on black 384-well or 96-well plates for optical imaging (Greiner) for Operetta CLS high-content confocal imaging. We used the non-confocal mode and a 10x air objective to image the well center in live-cell imaging mode with temperature and CO_2_ control at 37°C in a 5% CO_2_. Optimal z-position was determined by performing z-stack analysis on control wells prior to whole-plate acquisition. LED power and detector exposure time were adjusted on induced and non-induced control wells. Images were generally taken at 24h and 48h after induction or in a longitudinal setting over 72h with image acquisition every 2-4h. Longitudinal image acquisition and SARS-CoV-2 infected experiments were performed on 96-well plates for optical imaging (Greiner) using the Incucyte live-cell analysis system with temperature and CO_2_ control at 37°C in a 5% CO_2_. Infected cells were generally imaged every 2h after initial infection up to 5-8 days.

### Image quantification

Following acquisition, quantification was performed using the Harmony software (Perkin Elmer) for Operetta CLS images: After filtering each channel (sliding parabola 10px), we used the Harmony-Software building blocks to identify fluorescent objects above background mCherry intensities. Fluorescent objects were filtered by applying a threshold for object size and mean intensities as well as number of objects were determined. As a proxy for cell viability and fitness, cell density area was calculated from filtered brightfield images. Data with all relevant parameters was exported as csv files and analysed using R Studio and GraphPad Prism. Mean fluorescence and viability score were always calculated from technical replicates (n=3 or 4) and normalized to induced but non-drug treated control wells.

Quantification of Incubyte images was performed using the Incucyte analysis software (Sartorius): Incucyte software identifies whole cell layer area and fluorescent area above background within the image area and calculates mean fluorescence intensity. Data with all relevant parameters was exported as csv files and analysed using GraphPad Prism. Mean fluorescent area was calculated in relation to cell layer area to account for cell toxicity. Integrated mCherry intensity was calculated by factoring mean fluorescent area with mean fluorescence intensity to gain a total fluorescence score. Incucyte experiments were performed with technical replicates (n=4). Statistical testing was done through 1-way ANOVA with multiple comparisons testing.

The trend lines in Figure 4e are based on second order polynomial regression fitting, implemented through the GraphPad Prism v9.4 using the ‘‘Interpolation’’ function.

### ACE2 enzyme activity assay

ACE2 enzymatic activity was assessed using an MCA peptide-based substrate assay (kind gift of M. Lebedin and K. de la Rosa, Max Delbrück Center, Berlin, Germany) releasing a fluorophore upon cleavage by ACE2. For this, ACE2-expressing 293T and A549 reporter cells were seeded at 20000 cells per well on a 96-well plate for optical imaging (Greiner) and incubated overnight at 37°C and 5% CO_2_. Mca-APK(Dnp) substrate (Enzo life sciences) was diluted 1:100 in Epelman buffer (TrisHCl 75mM pH 6.5, 1M NaCl) supplemented with protease inhibitor mix and 100uM ZnCl_2_. Medium was removed prior to substrate addition. 20ul of diluted MCA substrate were added per well and fluorescence (Ex 340nm, Em 420nm) was measured every 30 min following substrate addition using the Tecan Spark (Tecan) with temperature control at 37°C. Activity was referenced to recombinant ACE2 protein and parental 293T and A549 reporter cells. Statistical testing was done through 1-way ANOVA with multiple comparisons testing.

### SARS-CoV-2 viral infection

All SARS-CoV-2 variants and associated experiments were handled in the biosafety level 3 (BSL-3) laboratory (L. Čičin-Šains laboratory, Helmholtz Centre for Infection Research (HZI), Braunschweig, Germany).

For 293T-ACE2-COVGT5 infection experiments, 20000 cells per well were seeded in 100ul of media and incubated overnight at 37°C and 5% CO_2_. Cells were infected with SARS-CoV-2 variants and subsequently treated with drug modulators at indicated concentrations. Longitudinal measurement of COVGT5 induction was performed as described above.

### Drug screening

High content drug screening for COVGT5 induction modulators was performed using the Operetta CLS (Perkin Elmer). For this, 293T, A549 and Calu-3 COVGT5 reporter cells were Seeded at 2000 cells per well (293T and A549) or 10000 cells per well (Calu-3) depending on cell growth rate in 50ul of respective media. Cells were kept at 37°C and 5% CO_2_ for 2 days until a confluency of 60-80% was reached. Reporter cells were treated with IFN mix or 3p-hpRNA (only 293T and A549) following the above-described procedure and incubated at 37°C and 5% CO_2_. 5-6h post initial induction, cells were treated with drug library. For this, drugs were prepared in a serial dilution setting using respective cell media to reach final concentration of 30nM, 300nM and 3uM in technical replicates (n=4) and final DMSO concentration <0.5%. Confocal fluorescence images were acquired 24h and 48h after initial induction following the above-described procedure. Image quantification was performed as described above. Data containing quantified parameters on COVGT5 fluorescence intensities, as well as cell area was imported as csv-files and analyzed using R (v.4.1.2) and R studio. After data merging and cleaning, drug concentrations were matched to COVGT5 intensities and cell area counts. For each cell line, timepoint and concentration, z-scores for COVGT5 mean intensity and mean cell area were calculated separately and merged back into a combined data frame. Hexagonal plots, bubble plots, spider plots and lollypop plots with COVGT5 and cell area z-scores were plotted using the ggplot2 (v.3.3.6) package. To generate circular heatmaps, the circlize package (v.0.4.15) was used.

### FACS analysis of reporter modulation

All analyses were performed using FlowJo_v10. For quantification of mean fluorescence intensity, mCherry positive gates were established following doublet exclusion. Mean fluorescence values of technical triplicates (n=3) for each treatment condition was calculated and normalized to IFN mix or 3p-hpRNA induced but non-drug treated condition. Data was exported as csv files and further analyzed using R studio and GraphPad Prism.

### Bliss drug synergy experiments

Drug and pathway synergy experiments were performed in 293T and A549 reporter cells using high throughput FACS readout. For this, cells were seeded on 96-well cell culture plates (Sarstedt) at 10,000 cells per well in 100µl of respective media. Cells were kept at 37°C and 5% CO_2_ for 2-3 days until a confluency of 60-80% was reached. First, cells were treated with IFN mix or 3p-hpRNA as described above and incubated at 37°C and 5% CO_2_ for 5-6h. Drug combinations were dispensed using the D300e digital dispenser (Tecan) in technical triplicates (n=3). For ionizing irradiation, cell plates were treated using the XenX irradiator (Xstrahl) under non-focused, open beam setting. Cells were incubated at 37°C and 5% CO_2_ for 2 days. FACS measurement and quantification was performed as described above. Data was exported as csv files and analyzed using GraphPad Prism to generate heatmaps and R studio to calculate bliss synergy using the synergyfinder (v.3.15) package. Mean synergy score was calculated through averaging across all combination datapoints. Statistical testing was done through 1-way ANOVA with multiple comparisons testing.

## Acknowledgments

We are grateful to E. von Weizsaecker (Ascenion) for connecting the Gargiulo and Cicin-Sain labs. We are also grateful to L. Li, H. Naumann, and the MDC FACS technology platform for technical support and to E. Wyler and M. Landthaler for timely sharing scRNA-seq data and A. Aguzzi for comments. BJ, CC, MJS are graduate students with Charité and Humboldt University. pscALPSpuro-HsACE2 (human) was a gift from Jeremy Luban (Addgene plasmid # 158081). The MCA peptide-based ACE2 enzyme activity assay was a kind gift from Mikhail Lebedin and Kathrin de la Rosa.

## Supplementary Materials for

**Fig. S1.**
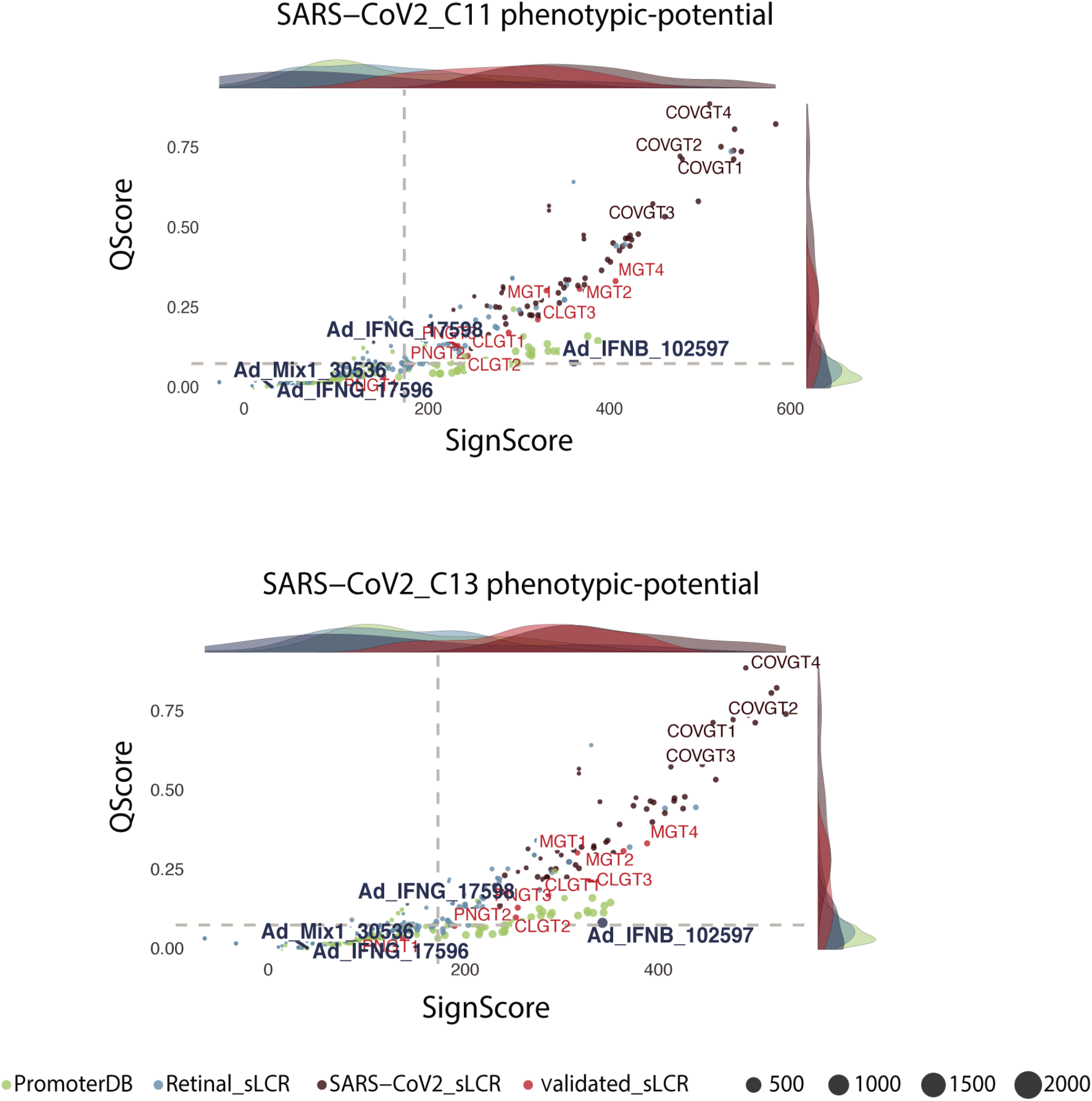
Activation potential of synthetic reporters in C11 and C13. Scatter plots showing signature (x-axis) and affinity (y-axis; methods) scores for the C11 (top) and C13 (bottom) phenotypes. Color codes separate reporters and sLCRs based on function and validation. Relevant signaling pathways of COVGT1-4 response. Relevant genes in C11 and C13 were assigned to respective signaling pathways and sorted by gene ratio and number of genes. Dot size indicates number of genes assumed to each pathway.

**Fig. S2.**
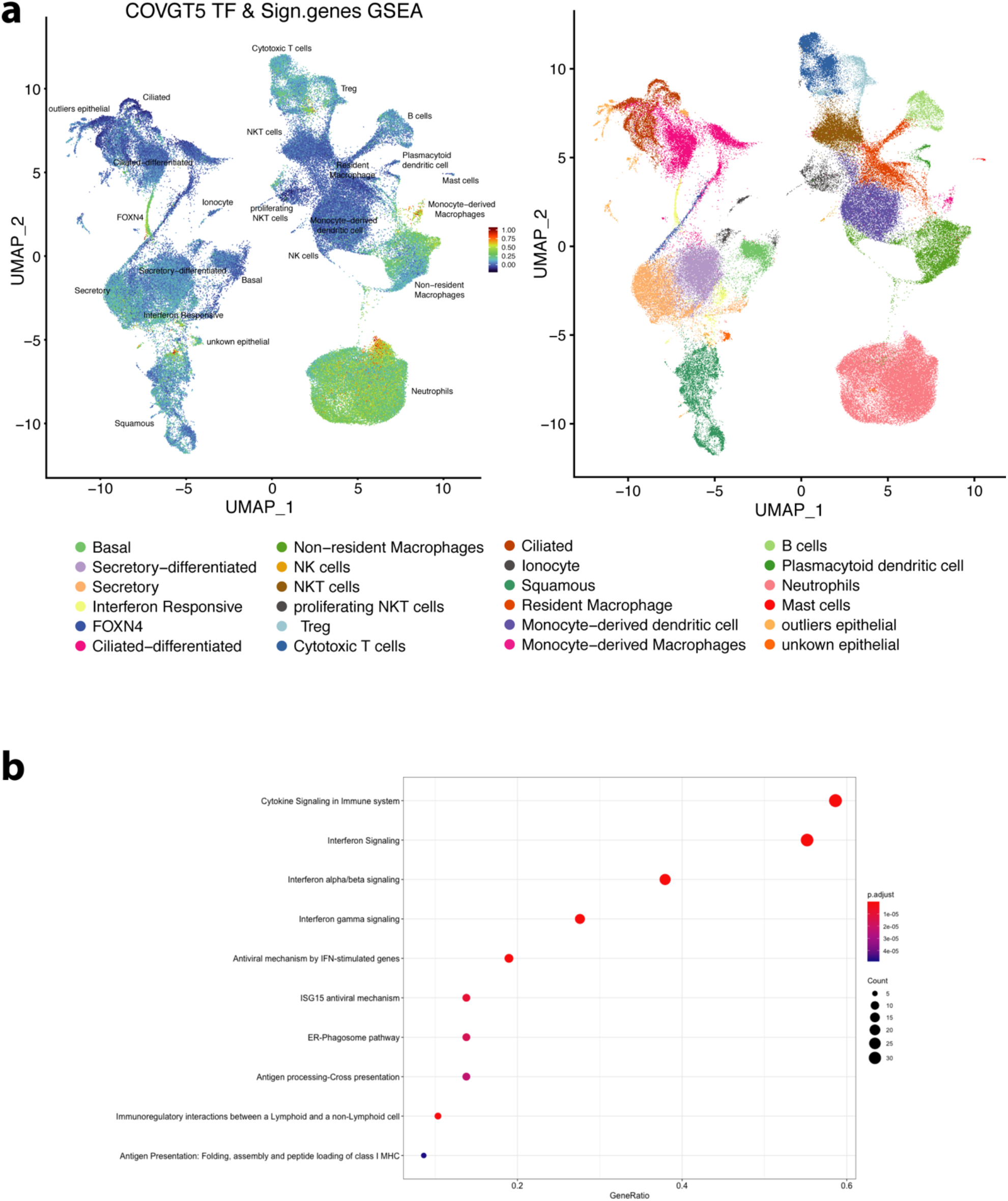
Representation of COVGT sLCRs in patient samples. **(A)** UMAP representation of COVGT5 gene coverage in SARS-CoV-2 patient samples. Gene set enrichment scores for COVGT5 signature genes and transcription factors on annotated cell clusters of Chua et al. 2020 are represented (left). Color code indicates cell populations marked (right). **(B)** Dot plot showing enrichment for signaling pathways of COVGT1-4 response. Signature genes common between epithelial cells and COVGT1-4 sLCRs were assigned to respective signaling pathways and sorted by gene ratio (X-axis value) and the number of genes (dot size).

**Fig. S3.**
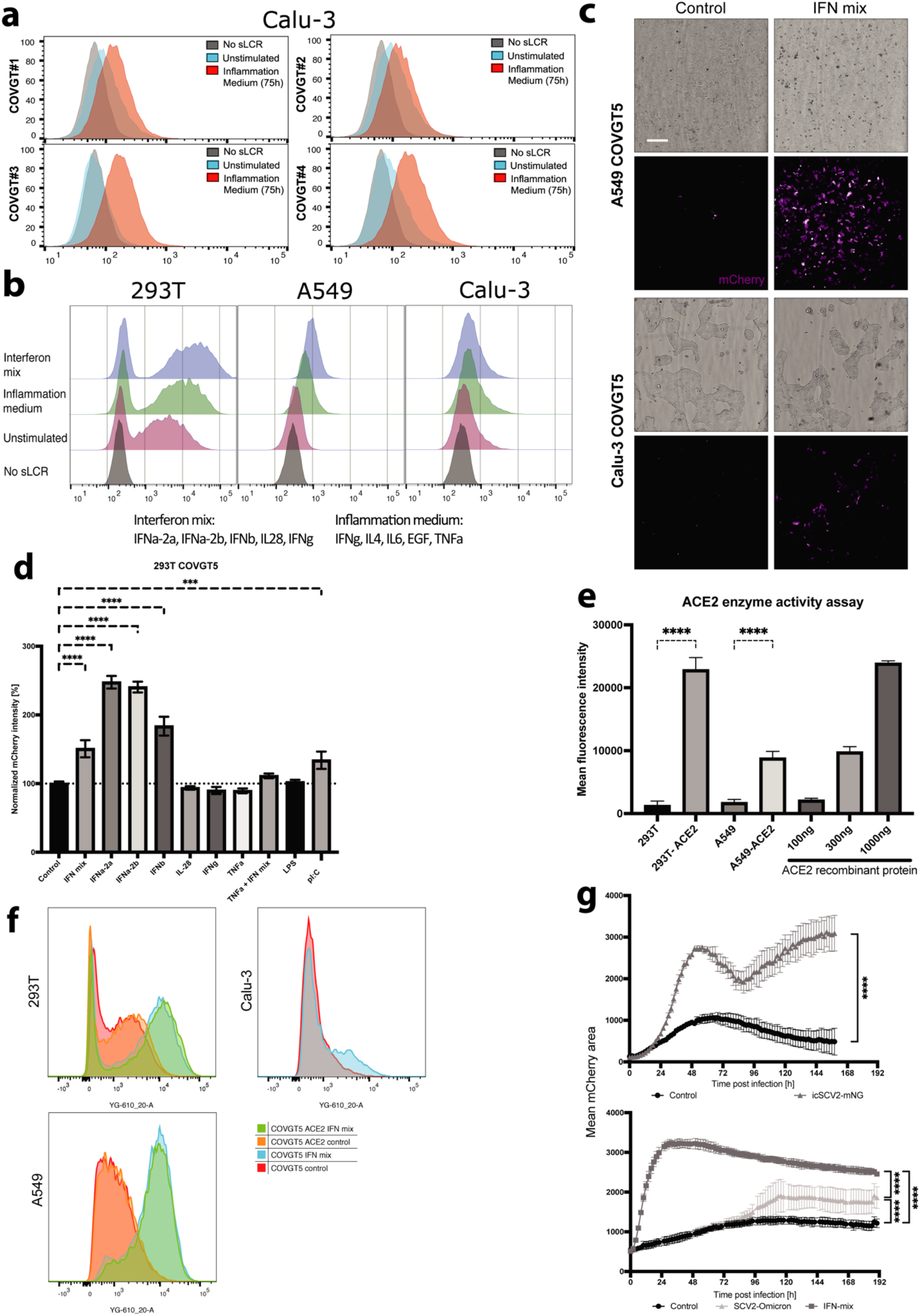
Further characterization of COVGT sLCRs response to triggers of epithelial immune response and SARS-CoV-2. **(A)**FACS analysis of Calu-3 with COVGT1-4 stimulated with inflammation medium for 75h **(B)**FACS analysis comparison between 293T, A549, and Calu-3 COVGT5 reporter cell lines. Cells were stimulated with ‘IFN’ or ‘inflammation’ mix for 48h. **(C)** Representative images of COVGT5-driven mCherry expression (purple) in A549 and Calu-3. Images were acquired by Operetta at 48h after IFN mix treatment. The scale indicates 200 um. **(D)** Bar plot quantification comparing induction of 293T-COVGT5 by individual cytokines, LPS, and pI:C. Reporter induction was assessed by normalization against untreated reporter cells 48h post-treatment. Mean mCherry intensity was calculated from technical triplicates (n=3). P-values denote significance by 1-way ANOVA. **(E)** ACE2 enzymatic activity of 293T and A549 ACE2 cells. Mean fluorescence was determined using an MCA-based peptide substrate assay comparing it to the activity of cell-free recombinant ACE2 protein. Fluorescence was quantified using the Tecan Spark in technical replicates (n=4). P-values denote significance by 1-way ANOVA. **(F)** FACS analysis comparing the inducibility of 293T and A549 reporter cells with their ACE2-modified variants. Cells were treated with the IFN mix and analyzed after 48h. **(G)** Longitudinal measurement of 293T-COVGT5 upon SCV2 infection. Cells were treated with 6600 pfu/ml icSCV2-mNG (top) or 88 pfu/ml SCV2-Omicron and IFN mix (bottom). Measurement was performed using the Incucyte. The relative mean fluorescent area was determined from technical replicates (n=4). P-values denote significance by 1-way ANOVA.

**Fig. S4.**
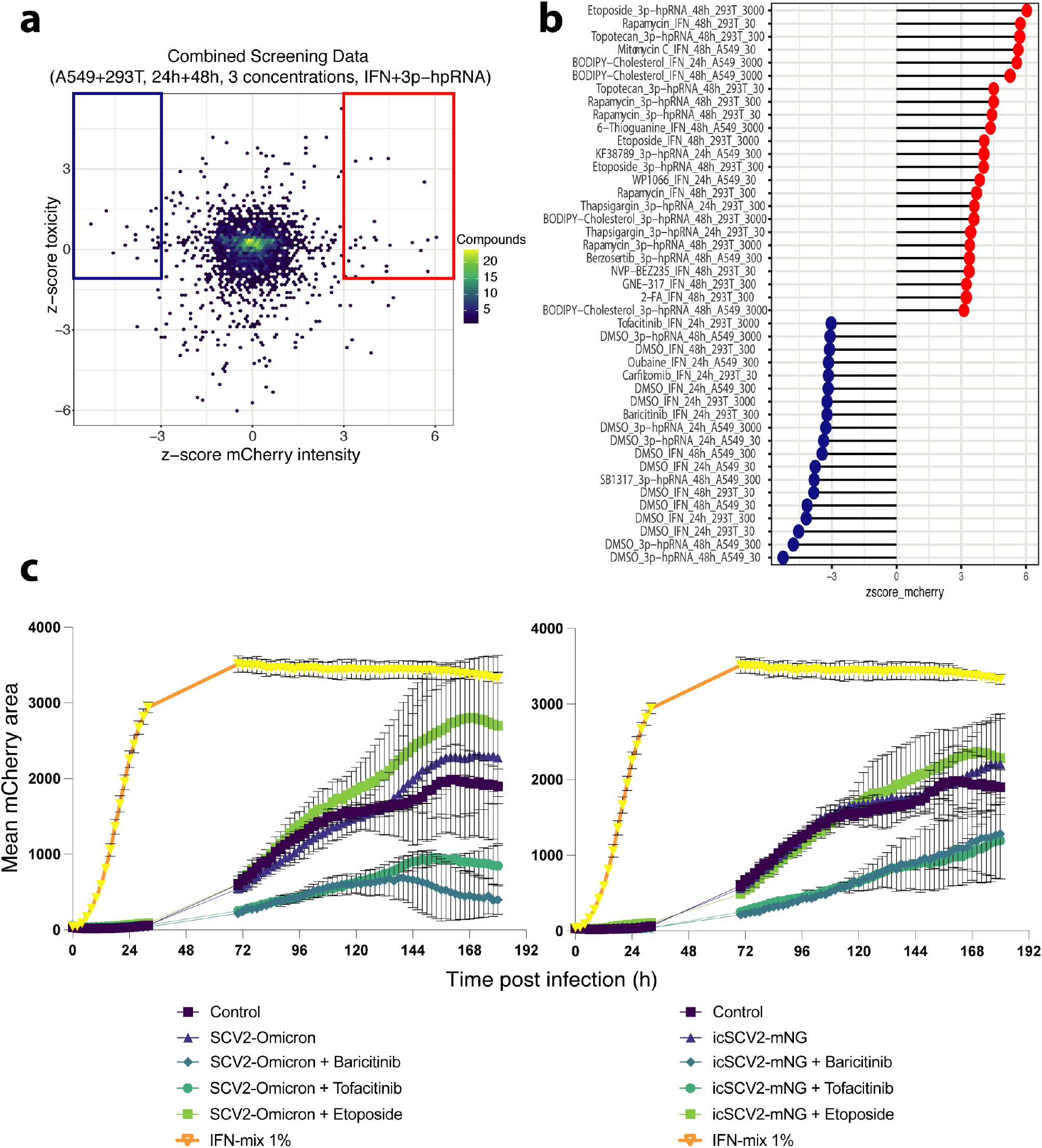
Alternative representation of COVGT5 screening results and SARS-CoV-2 validation experiments. **(A)** Combined representation of 293T and A549 screening results. Z-scores for Toxicity (y-axis) and mCherry intensity (x-axis) are plotted against each other. Marked areas indicate drugs with no negative effect on cell viability and amplifying (red) or inhibitory (blue) effect on COVGT5 induction. The color scale indicates the number of compounds in each hexagon. **(B)** Lollipop plot showing marked data points from a) ranked based on COVGT5 z-scores. **(C)** Longitudinal measurement of 293T-COVGT5 upon SCV2 infection. 6600 pfu/ml SCV2-Omicron (left) or icSCV2-mNG (right) were used. Measurement was performed by Incucyte. The relative mean fluorescent area was determined from technical replicates (n=4).

**Fig. S5.**
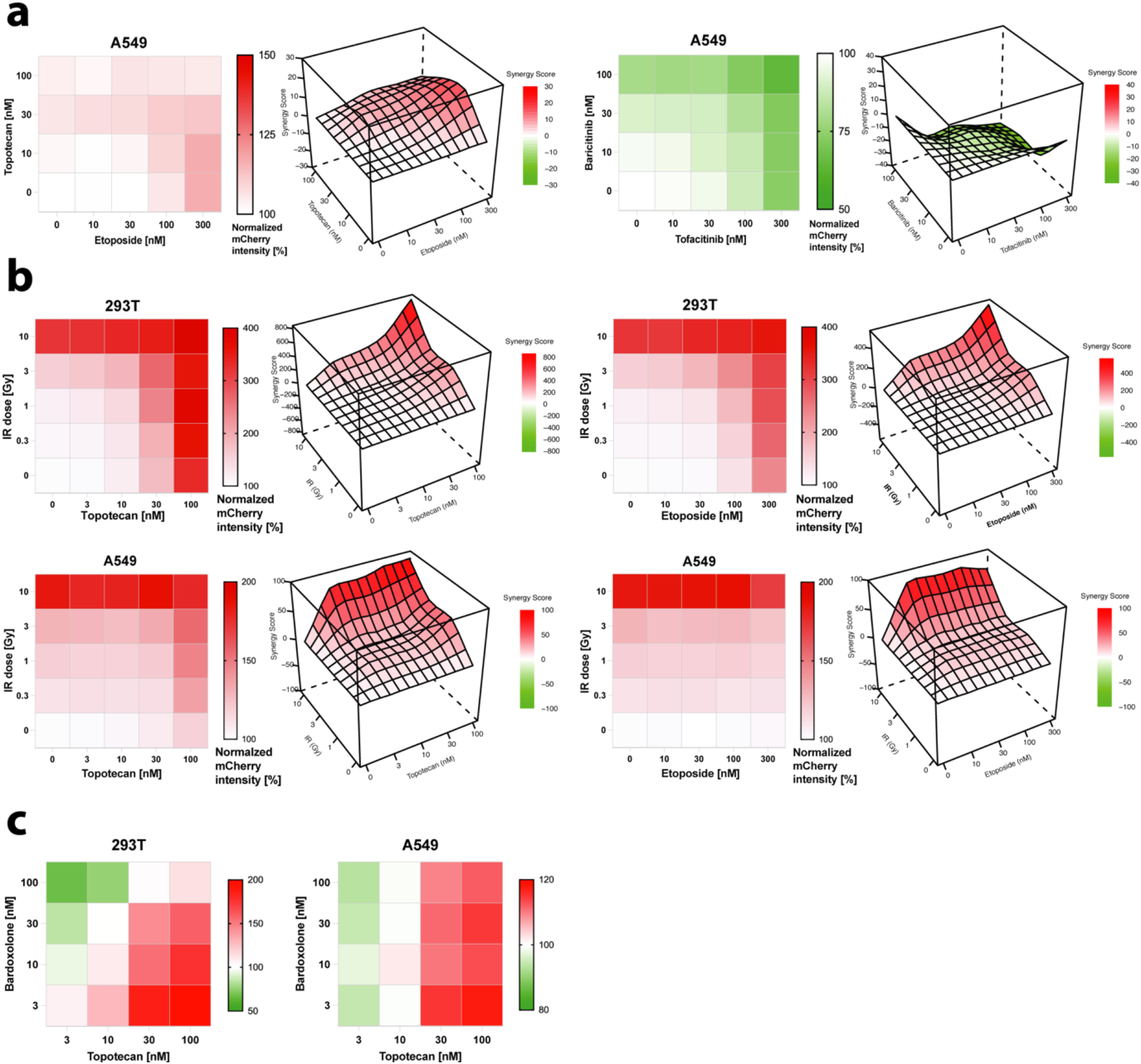
Extended data on synergistic regulation of COVGT5. COVGT5 reporter induction and mCherry quantifications were acquired using high throughput FACS by measuring mean mCherry intensity of technical triplicates (n=3) and normalized to non-drug-treated cells. **(A)** Dose-escalation of Topotecan against Etoposide (left) and Baricitinib against Tofacitinib (right). A549-COVGT5 received IFN mix induction followed by combinatorial drug treatment. Bliss synergy scores inform on synergism between Topotecan + Etoposide (left) and Baricitinib + Tofacitinib (right) in modulating COVGT5 response. **(B)** Dose-escalation of IR against Topotecan (left) and against Etoposide (right). 293T (top) and A549 (bottom) reporter cells received 0.1 ug/ml 3p-hpRNA transfection followed by IR and drug treatment. Bliss synergy scores inform on synergism between IR + Topotecan (left) and IR + Etoposide (right) in modulating COVGT5 response. **(C)** Dose-escalation of Bardoxolone against Topotecan in 293T (left) and A549 (right) reporter cells. Type or paste caption here. Create a page break and paste in the Figure above the caption.

**Table S1.**

Table S1 is provided as a separate file.

**Table S2.**

Table S2 is provided as a separate file.

